# Disease Network Delineates the Disease Progression Profile of Cardiovascular Diseases

**DOI:** 10.1101/2020.09.09.290585

**Authors:** Zefang Tang, Yiqin Yu, Kenney Ng, Daby Sow, Jianying Hu, Jing Mei

## Abstract

As Electronic Health Records (EHR) data accumulated explosively in recent years, the tremendous amount of patient clinical data provided opportunities to discover real world evidence. In this study, a graphical disease network, named progressive cardiovascular disease network (progCDN), was built based on EHR data from 14.3 million patients ^1^ to delineate the progression profiles of cardiovascular diseases (CVD). The network depicted the dominant diseases in CVD development, such as the heart failure and coronary arteriosclerosis. Novel progression relationships were also discovered, such as the progression path from long QT syndrome to major depression. In addition, three age-group progCDNs identified a series of age-associated disease progression paths and important successor diseases with age bias. Furthermore, we extracted a list of salient features to build a series of disease risk models based on the progression pairs in the disease network. The progCDN network can be further used to validate or explore novel disease relationships in real world data. Features with sufficient abundance and high correlation can be widely applied to train disease risk models when using EHR data.

## Introduction

Accumulating evidence have shown that diseases are associated with each other^1–5^. Since molecular components were functionally interdependent in a human cell, a disease was rarely caused by abnormality of one single gene but the perturbations of the complex intracellular network^6^. In other words, diseases were associated due to the intersection of their underlying molecular components. For example, the action potential of a cardiomyocyte required the coordinated actions of more than twenty different ion transporters and channels, leading to complex interactions underlying cardiovascular diseases such as atherosclerosis, cardiac hypertrophy, heart failure and arrhythmias^1^. Metabolic syndromes, such as glucose intolerance, obesity, hypertension, and dyslipidaemia,^5, 7^ also showed dysregulation associations in essential components.

Disease network is a well-known approach to delineate the relationships among human diseases^2, 4^. A seminal work, the human disease network (HDN)^2^, leveraged the gene mutation and phenotype data to discover disease connections. The edge between two disease nodes represented that at least one gene mutation was shared between these two diseases. While HDN discerned the genetic origins or general patterns of human diseases, other studies^8, 9^ identified disease-disease associations based on similarity of mRNA, microRNA expression for predicting new uses of existing drugs, namely, drug repurposing.

However, estimating the disease connection solely based on the molecular level could be misleading since the same mutation or similar expression may not determine disorders in real clinical diagnoses. Real world data has the potential to address this challenge because it captures the actual clinical observations. With the explosive accumulation of electronic health records (EHR) data, building disease networks based on real world data became a reality. Recent studies identified the co-occurring diagnoses among diseases as disease connections^10–12^. The identified highly correlated diseases or disease modules had biological interpretations, which provided evidence that this could be a new strategy for driving further understanding of diseases.

Although existing disease networks are able to identify a series of disease connections, they do not use the temporal information to identify disease progression profiles. In order to overcome this challenge and identify the temporal associations among human diseases, we propose the progressive cardiovascular disease network (progCDN) to discover the disease connections and temporal associations among cardiovascular diseases. The EHR records from 14.3 million cardiovascular disease (CVD) patients were collected for network construction. Although several previous studies had constructed disease networks on CVDs^1, 13–16^, most of them focused only on undirected associations among diseases, such as whether they share common genes. In our study, we leverage the time stamps of diagnoses to build progCDN with directed edges based on temporal associations. These temporal patterns extracted from the large data set provided us with opportunities to identify novel disease connections.

We used new designed methods, progression rates (Methods), to delineate a comprehensive progression profile of diverse CVDs in progCDN. Using the network, we observed general and specific disease progression paths among CVD patients. An imbalanced distributions of disease progression profile was also observed in different age groups. According to the disease network, we identified EHR data features with high performances and developed a series of risk models.

## Results

### Properties of disease network

Using the disease progression associations identified from EHR data, we built the progressive cardiovascular disease network (progCDN) to capture the progression and comorbidities of CVDs (Figure 1). The network contained 93 SNOMED CT-based diseases^17^ as nodes, including 35 source nodes (predecessors) and their target nodes (successors), with 796 directed edges (Methods, Table 1). The nodes were classified into 9 disease classes and labeled with corresponding colors. The directed edges (with arrow) indicate the progression path from predecessor to successor. We observed that except for 35 selected predecessors, the other 58 diseases in different classes, such as hematologic, nutritional, genitourinary and infectious diseases, were diagnosed among the patients who had CVDs. Meanwhile, the patients with CVDs also had a tendency to develop more serious CVDs, such as coronary arteriosclerosis or heart failure, which indicated the complexity of the CVD progression pattern.

**Figure 1:**
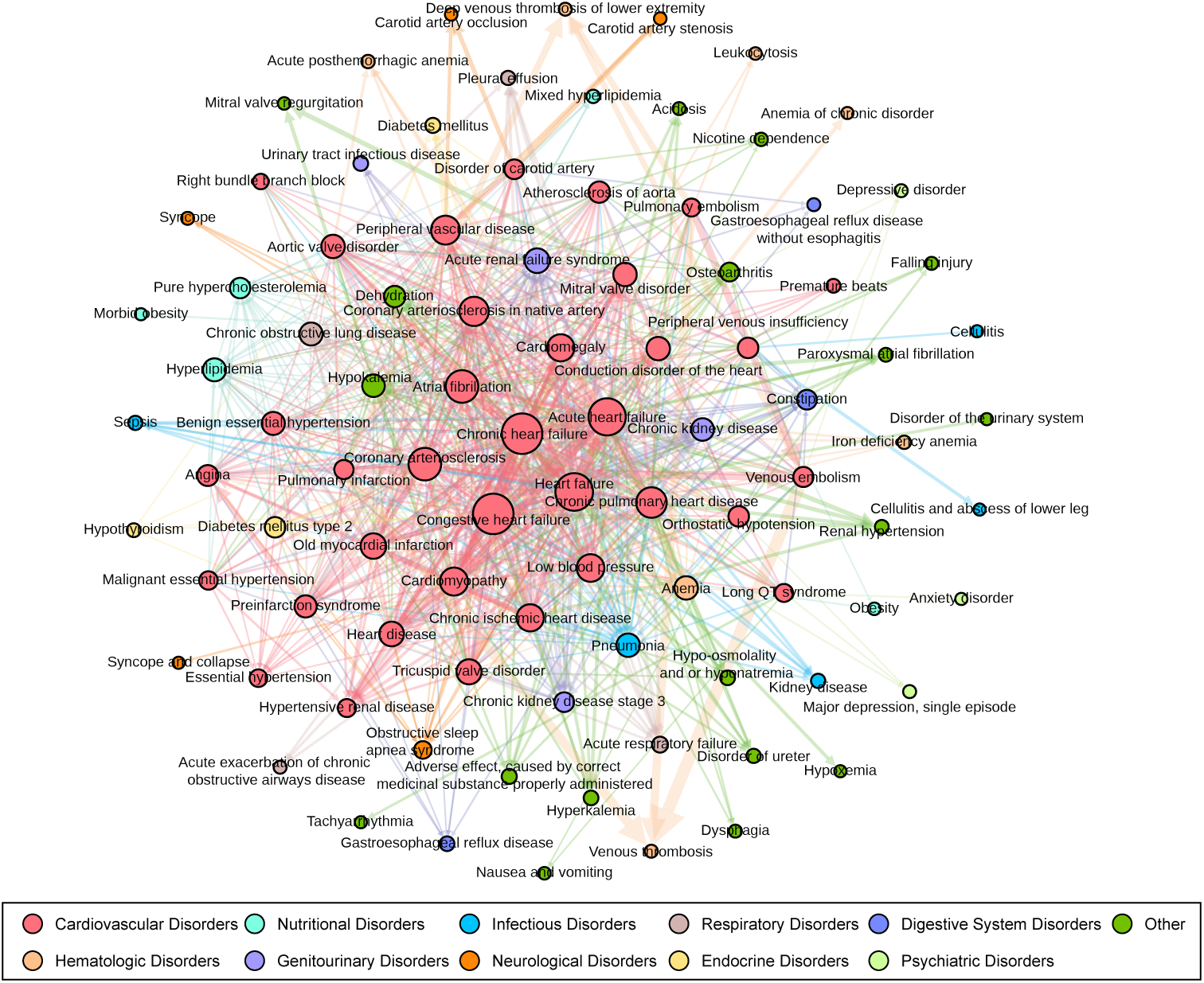
Disease progression network of the CVD diseases. Nodes represent diseases based on Snomed-CT system. Each node was classified into different disease clusters with corresponding color (red for cardiovas-cular diseases, light blue for nutritional diseases, blue for infectious diseases, brown for respiratory diseases, light orange for hematologic diseases, violet for genitourinary diseases, violet for genitourinary diseases, orange for neurological diseases, yellow for endocrine diseases and green for other diseases). The size of the node represents the degree value of the disease. The edges with array indicate the progression path from source node (predecessors) to target node (successors). The color of edge is identified by the target node class.

Next, we focused on the nodes with the highest number of direct connections (Figure 2A). By ranking the number of connections (degree property D), we observed that heart failure (including congestive heart failure [D = 69], chronic heart failure [D = 69], heart failure [D = 62] and acute heart failure [D = 61]), coronary arteriosclerosis (D = 50) and atrial fibrillation (D = 50) had the most direct links with other diseases, indicating that these diseases were dominant in disease development and these diseases should be managed with high priority in medication. On the other end of the scale, several diseases, such as tachyarrhythmia, Hypoxemia and cellulitis, had only one direct connection with other diseases.

**Figure 2:**
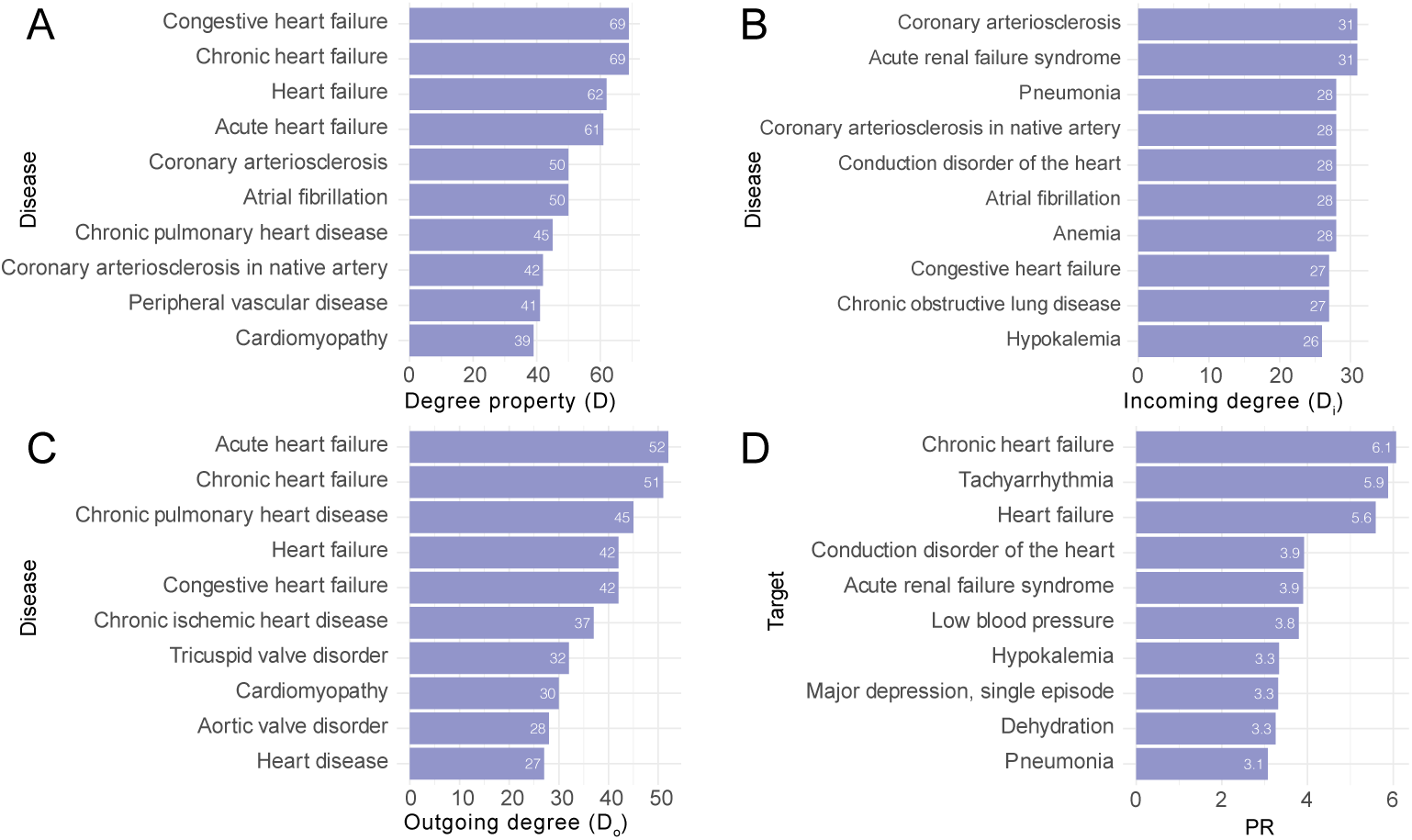
Bar plots indicate the degree of disease network and the progressed diseases of long QT syndrome. The top 10 nodes with highest degree (A), incoming degree (B) and outgoing degree (C) of the disease network. (D) The top 10 successors with highest progression rate of long QT syndrome.

To discover the important predecessors and successors, we ordered the nodes by outgoing degree (Do) and incoming degree (Di) (Figure 2B, C). The acute heart failure (Do = 52) and chronic heart failure (Do = 51) nodes had the highest number of outgoing connections, indicating that these diseases are likely to progress to more severe conditions. Coronary arteriosclerosis and acute renal failure syndrome (Di = 31) have the highest number of incoming connections, indicating that there are many diverse pathways leading to these conditions and these diseases should be concerned in CVD medication. Though a lot of important successors are CVDs, we also found that CVD patients are likely to develop pneumonia and sepsis (infectious disorders), acute respiratory failure (respiratory disorders), or chronic kidney disease (genitourinary disorders), which are consistent with previous studies^18–22^ (Supplementary Table 1). Additionally, progCDN also delineated several specific progression pairs, such as peripheral venous insufficiency followed by cellulitis and hypoxemia followed by heart failure.

Since our EHR cohort contained a very large volume of patient records, we further interrogated the progression paths of the rare disease, long QT syndrome (LQTS) (Figure 2D), which is a heart rhythm condition and is identified by abnormal interval prolongation on the electrocardiogram. This disorder could cause fast, chaotic heartbeats and even sudden cardiac death in young individuals with normal cardiac morphology^23^.

In our data, we observed that LQTS happened among young people (Supplementary Figure 1). After the diagnosis of LQTS, patients are more likely to develop chronic heart failure, tachyarrhythmia and heart failure, which is consistent with current knowledge^24, 25^. Notably, major depression is also a successor of LQTS (PR = 3.3). However, few publications studied this relationship^26^, which requires further studies.

### Disease progression across age groups

Though CVDs are diagnosed in all ages (Supplementary Figure 1), patients in different age groups may have different disease progression patterns. To investigate the disease developments across the strata of age, we divided patients into 3 age groups, young (age from 18 to 35), middle-aged (age from 36 to 55) and elderly (age from 56 to 90) referring to previous strategy^27^.

Three progCDN instances are constructed according to the different age groups (Supplementary Figure 2-4). We found 59 common successors were shared among these age groups (Figure 3A) and 130 successors were specific to each age group (20 for young, 24 for middle-aged, and 86 for elderly). In addition, the middle-aged group shared 57 diseases with the elderly group, which were far more than the successors shared between the young and middle-aged groups (15 diseases) as well as the ones shared between the young and the elderly disease groups (3 disease), indicating that patients age 36 have more commonalities.

**Figure 3:**
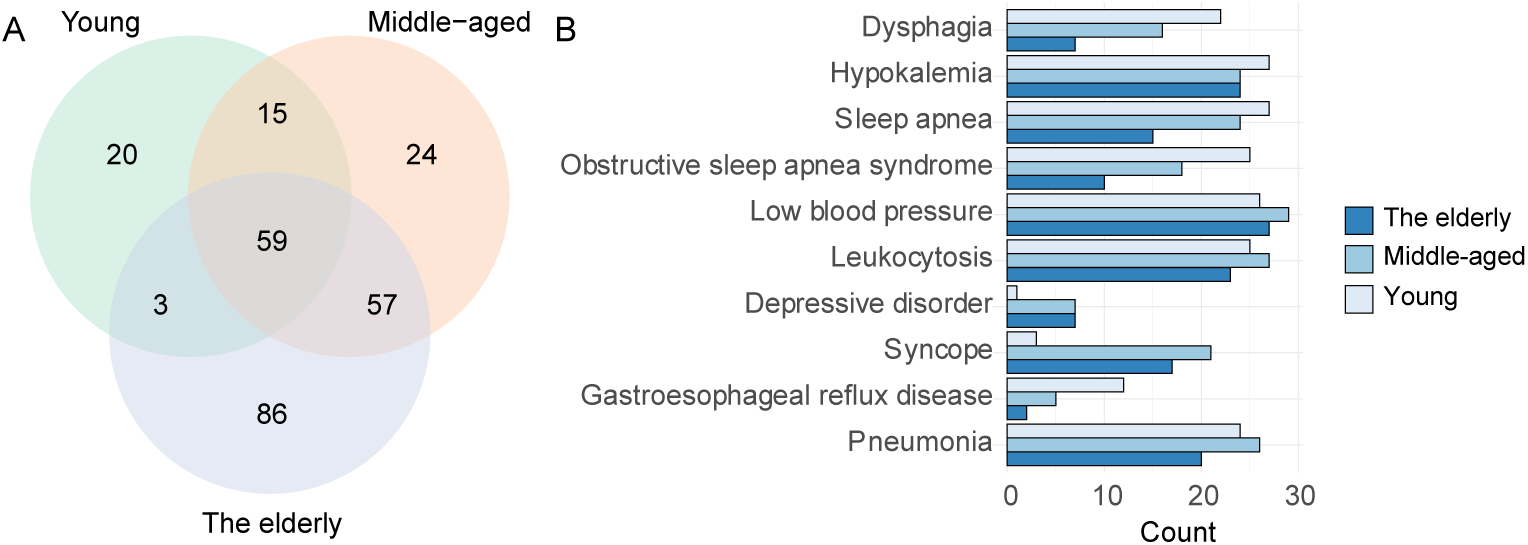
The distribution of progressed diseases among young, middle-aged and the elderly disease networks. (A) Venn plot described the shared and unique successor diseases among different age group. (B) The count distribution of shared successor diseases in young (light blue), middle-aged (blue) and the elderly (dark blue) disease networks.

Among 59 common successors, some were general conditions, such as low blood pressure, hypokalemia and anemia (Supplementary Table 2), which have been previously reported as the comorbidities of cardiovascular diseases^28–31^. A series of diseases showed imbalanced distribution in different age groups (Figure 3B). For example, patients in the middle-aged and the elderly groups tend to have syncope after the diagnosis of CVDs, while dysphagia and obstructive sleep apnea syndrome are more likely to occur after CVD diagnosis in the young CVD patient group. Although syncope and obstructive sleep apnea syndrome are both highly associated with cardiovascular diseases, the age distribution of them have not been discussed before^32^. In addition, the relationship of dysphagia and CVDs was seldomly reported before, which is another novel finding to be studied in the future.

Further, we focused on the degree property (D) and the incoming degree (Di) to identify important nodes in different age groups. Heart failure (as well as chronic heart failure, congestive heart failure and acute heart failure) occurred in all age groups with a high rank of degree property, which are consistent to the general disease network (Figure 2A). When ranking the incoming degree of nodes in each age group, we observed that hypokalemia, sleep apnea and pneumonia are more likely to be a successor condition in the young group. Serious diseases, such as old myocardial infarction, acute renal failure syndrome, hyperkalemia and chronic kidney disease, performed as important successor diseases in the middle-aged group with high incoming degree value. In addition, the severe diseases, such as chronic diastolic heart failure, tricuspid valve disorder and chronic pulmonary heart disease are more likely to be targeted in the elderly group.

Additionally, we also found several age-associated disease pairs with high PR values (Supplementary Table 1). In the young group, patients who had acute heart failure would progress to low blood pressure (PR = 10.8), which is consistent to previous studies^33^. The cardiomyopathy patients in the middle-aged group are likely to have heart failure in the next few years (PR = 25.1) while pulmonary infarction (PR = 25.7) and pulmonary embolism (PR = 22.9) tends to progress to venous thrombosis in the elderly group.

### Disease network identifies risk models for disease prediction

ProgCDN indicated the connections among diseases, which can help us understand the progression of CVDs. Risk models based on these progression pairs could benefit patient in clinical treatment. However, the traditional strategy for designing risk models requires a long time in collecting patient cohort and expert-advised features. Further, the expert-advised features sometimes are unavailable or difficult to obtain in EHR databases. Thus, it is beneficial to identify features that are highly correlated with successors of interest and sufficiently available in EHR databases. The IBM Explorys database^34^ contains rich medical test data and demographic information, which enabled us to identify important features. Using the identified features, a series of risk models could be constructed in a short time. Hence, we next used a data-driven strategy (Figure 4A) to select the most correlated features of given disease pairs and used these features to build risk models.

**Figure 4:**
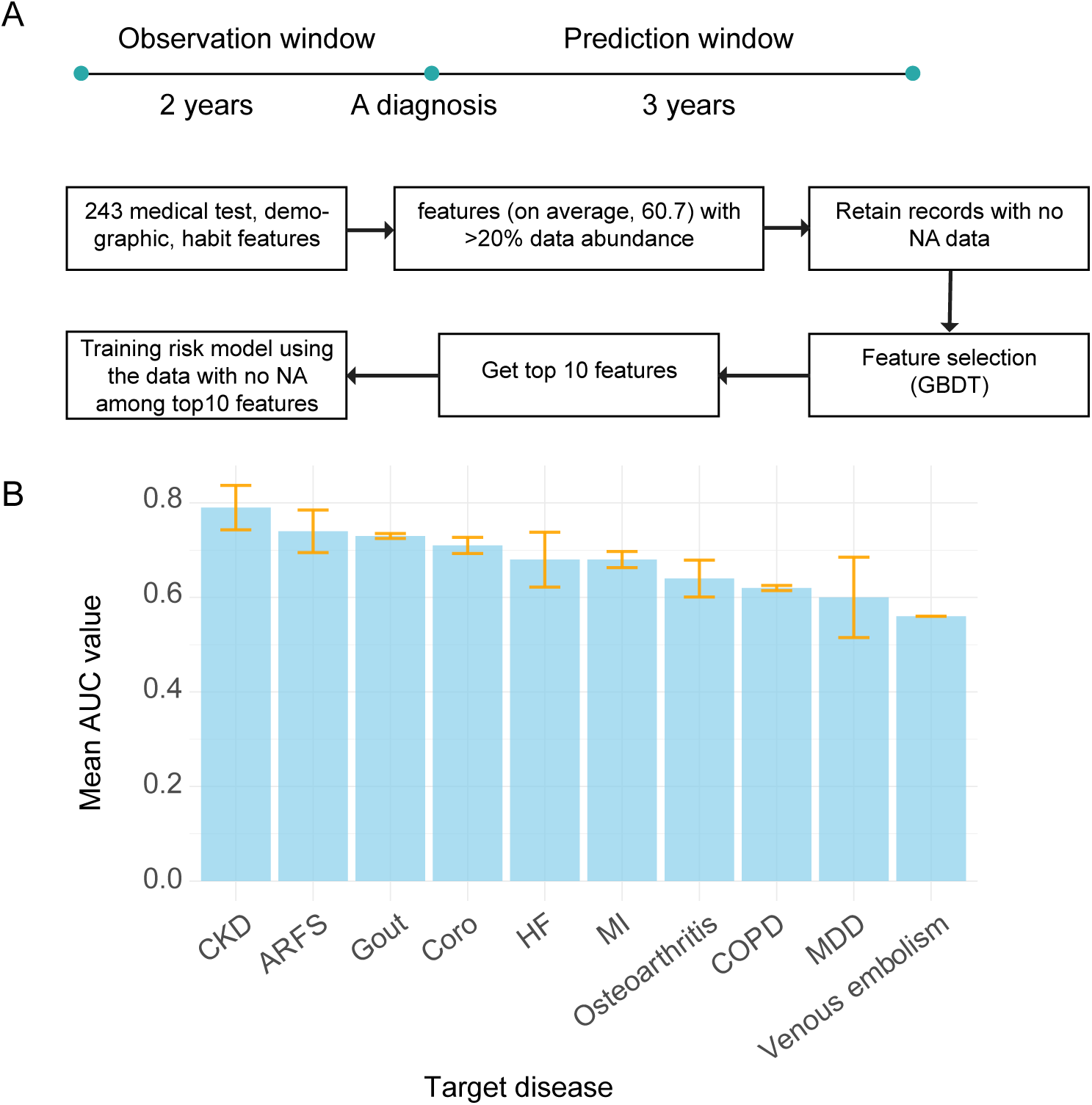
Identification of important features in EHR database. (A) The schema presents our strategy to identify top 10 features of given disease pair and build corresponding risk models. (B) The mean AUC values of the risk models grouped by target diseases.

We selected 23 disease pairs (Supplementary Table 3) based on ProgCDN for feature selection and risk model development. By extracting the features with sufficient data in the EHR database, we collected 243 features as a preliminary dataset for feature selection (Methods). As the pipeline showed (Figure 4A), for each disease pair, the top 10 features with highest importance value were retained (Supplementary Table 4).

We observed that identified features were highly associated with the successors. For example, urea nitrogen and creatinine are identified as key features in the risk model for the hyperlipidemia-CKD (Chronic kidney disease) pair, which is consistent to the knowledge that these two features are usually used for glomerular filtration rate estimation (GFR)^35^. Though the GFR and cystatin C are important biomarkers for kidney disease^36^, these two features are filtered in our pipeline due to poor data abundance, indicating that parts of well-known features may not have sufficient data abundance in EHR data. Meanwhile, we also detected several important but rarely reported features, such as thyrotropin and hematocrit, which could be regarded as candidate biomarkers in the future studies.

By using the top 10 features of each disease pair, we next built a series of risk models with 2-year observation window and 3-year prediction window (Methods). By ranking the AUC value of risk models, we observed that some risk models had excellent performance (Supplementary Table 4), such as Type 2 diabetes mellitus (T2DM) to CKD (AUC = 0.83) and Hyperlipidemia to CKD (AUC = 0.83). However, a few risk models, such as AF to Venous embolism (AUC = 0.56) and LQTS to major depression disorder (MDD) (AUC = 0.54), showed bad performance, which indicated that not all disease progression pairs were appropriate for designing risk models in our EHR database. Since the disease pairs with the same successors showed similar performance, we next grouped the risk models by successors (Figure 4B). We observed that kidney diseases, such as chronic kidney diseases and acute renal failure syndrome (ARFS), performed much better than psychological diseases such as MDD. This situation may be caused by the features extracted from the EHR database. The majority of the data in the EHR database are laboratory test results, which could represent the metabolic status but cannot reflect the mental situation of patients. Thus, the kidney diseases, which could be predicted by biomarkers, would have better performance.

Further, to identify the frequently used features among these features, we collected the features in the high-performance risk models (AUC > 0.65) and calculated their mean rank level. Urea nitrogen, platelet mean volume, age, creatinine and albumin are the top 5 ranked features, which are used frequently in our selected risk models (Supplementary Figure 5). These frequently used features are correlated with the diseases and have sufficient data in the EHR database, which could provide guidance for further research of prediction models based on EHR data.

## Methods

### Disease network construction

To analyze the progression pattern and comorbidities of cardiovascular diseases, we collected the patients with cardiovascular disease diagnoses from the IBM Explorys database^34^ to construct the cohort. The cohort contained 14.3 million patients aged 18 to 90 years old who have diagnosis information between 2010-2016. Next, 31 CVD diseases with high frequency of diagnosis (frequently used SNOMED CT items^17^) were selected as predecessors in the disease network. Hypertension, hyperlipidemia and diabetes mellitus type 2 are known to be highly associated with CVD diseases and were also included as predecessors. Since the sample size was large enough, a rare CVD disease, Long QT syndrome, was also listed as a predecessor for further analysis (Table 1).

In the disease network, each node represents a disease with an associated SNOMED CT code^17^. Two nodes are connected by a directed link if one has progression tendency towards the other one. To identify the progression tendency, we designed a new measurement, progression rate (PR), to measure the correlation degree between two diseases.

The nodes and edges with *PR >* 2.5, *I*_random_ *>* 0.5% were retained for disease map construction (Figure 1; Supplementary Figure 2-4). For visualization of Figure 1, we kept the nodes and edges with *I*_observed_ *>* 5%. The network software Gephi^37^ and Fruchterman Reingold clustering algorithm^38^ were used to construct the disease network. The nodes were clustered into major disease classes according to previous work^39^.

### Feature selection

To investigate the important features for developing risk models, we empirically selected 23 reasonable disease pairs in the disease network. 239 medical lab test features (Table 2) with sufficient data abundance (at least 1,000,000 patients records), age, gender, habits of alcohol and tobacco use were collected for feature selection. For each disease pair, we set a 2-year observation window before the diagnosis of the predecessor diseases and a 3-year prediction window after. In the observation window, the median value of all laboratory tests during these 2 years were computed. Outcome was set as 1 if the successor disease happens (or 0 if the successor disease doesn’t happen) within 3 years after the predecessor disease was diagnosed.

Thus, for each disease pair, we used a dataset with 243 features for feature selection. To extract the features with sufficient data towards a given disease pair, we filtered the features whose data abundances were less than 20% among these 243 features (on average, 60.7 features retained among the different disease pairs). To avoid losing information, we filtered the records with any drop-out among all features for feature selection. GBDT (Gradient Boost Decision Tree) algorithm was next used for calculating the feature importance. Further, the top 10 features ranked by importance values were kept as the most important features of a given disease pair.

After feature selection, the patient records with the top 10 features were collected to build risk models. Records without any missing data were kept for training and testing. We used 3 common binary classification algorithms, LR (Logistic Regression), RF (Random Forest) and GBDT models, to build the risk models. The best AUC value among these 3 algorithms was used as the prediction performance.

## Discussion

The analysis of EHR data enabled us to reveal real-world evidence (RWE) including the discovery of temporal disease relationships. The comorbidities and progressed diseases of CVDs identified from RWE can provide guidance for prevention and treatment in clinical practice. By analyzing 14.3 million patients records, which is the largest cohort so far, we built a CVD progressive disease network (progCDN) that reflects the real-world connections of among these diseases. We discovered common progressed diseases of CVDs, such as sepsis and acute respiratory failure, as well as several specific progression paths, such as cellulitis from peripheral venous insufficiency. The networks across the strata of age showed different progression patterns in different age groups. For example, we identified common progressed diseases with imbalanced age distribution, such as syncope, dysphagia, obstructive sleep apnea syndrome and osteoarthritis, as well as age-group specific diseases such as acute heart failure to pneumonia in young patients. Finally, a series of features with high correlation and abundance were extracted, which can be used by researchers to train risk models using EHR database.

Although our progression rate approach was designed to reveal the sequential diagnoses after a diagnosis of CVD, not all identified diseases exhibited an expected disease progression pattern. For example, patients tend to be diagnosed as “coronary arteriosclerosis in native artery” after the diagnosis of “heart failure” (PR = 7.6). However, based on common knowledge, “heart failure” is more serious compared with “coronary arteriosclerosis in native artery”. This behavior may be explained by the following reasons. First, a patient who already had “coronary arteriosclerosis in native artery” might not get the diagnosis due to no admission. Then, when he was admitted to the hospital because of the heart failure, the coronary arteriosclerosis in native artery would be diagnosed and recorded in EHR. Second, some of the diagnoses were used for supplementary medication, such as gastroesophageal reflux disease. For example, the traditional CVD drug aspirin is reported to cause the damage of gastrointestinal mucosa in the upper and lower gastrointestinal tract. The proton pump inhibitor (PPI), a medication for gastroesophageal reflux disease, could be used as combination therapy to prevent the damage^40, 41^. To get the medication of PPI, patients would be diagnosed as gastroesophageal reflux disease, which indicated that the gastroesophageal reflux disease we observed in the successor list might not be the real progressed disease. Thus, although our novel algorithm is designed to identify progressed diseases, the results still include many comorbidities, which implied that the diagnosis information in the EHR data usually do not obey the time sequence strictly.

The progression rate is a new measurement for representing the degree of progression relationship. According to its definition, the progression rate is similar to odds ratio (OR), which shows that the disease pair with PR > 1 has a progression relationship. However, we found that parts of the PR values of randomly selected disease pairs were also greater than 1. It may be partially explained by the enrichment of diagnoses among admissions. Hence, this situation required us to set a stricter threshold for identifying the progressed diseases. Hence, we used permutation strategy (Supplementary Figure 6; Supplementary materials) to get a suitable threshold PR = 2.5 for further analysis.

The disease network enabled us to identify a series of potential risk models. For kidney disease, heart failure, coronary arteriosclerosis or gout, the important features we identified, such as erythrocyte distribution width, urea nitrogen, albumin, intravascular systolic, and creatinine, are highly correlated with them. However, diseases such as major depression disorder or venous embolism may not be reflected by regular laboratory test results. Indeed, for mental disease, researchers used the features such as Hamilton Depression Rating Scale^42^ or metabolite profile^43^ to predict major depression. These features are not available in general EHR databases. Thus, although the data in EHR data contain thousands of lab test features and many patient records, these data are not suitable for predicting all diseases.

Furthermore, in our disease network, we use SNOMED CT items acting as nodes. Since the definitions of diseases in SNOMED CT are clear and meticulous, we observed the connections at a high resolution. Though the experts suggested that we need to merge parts of diseases together, such as heart failure, acute heart failure, chronic heart failure and congestive heart failure, we still use the intrinsic SNOMED CT items as nodes for delineating comprehensive real-world evidence.

In conclusion, the progCDN exhibited the disease progression profile among CVDs, which showed a series of well-known progression pairs as well as novel progression paths. Future studies could focus on the hypotheses of causal relationship based on the pairs identified from progCDN. The identified features with high correlation and abundance occurrence provided guidance for experts or researchers to build models using EHR data. This work provides new avenues to leverage real-world data and new strategies to analyze them, which partially unleash the treasure of these precious data.

## Footnotes

1 Data of 14.3 million individuals, pooled from multiple different healthcare systems with distinct EHR, were obtained. Data were standardized and normalized using common ontologies, searchable through a HIPAA-enabled, de-identified dataset (IBM Inc.). Patients were cardiovascular patients cohort seen in multiple healthcare systems from 2010 - 2019 with a combination of data from clinical EMRs, healthcare system outgoing bills, and adjudicated payor claims.

## References

1. Aldons J. Lusis and James N. Weiss. Cardiovascular Networks: Systems-Based Approaches to Cardiovascular Disease. Circulation, 121(1):157–170, January 2010. 368.

2. K.-I. Goh, M. E. Cusick, D. Valle, B. Childs, M. Vidal, and A.-L. Barabasi. The human disease network. Proceedings of the National Academy of Sciences, 104(21):8685–8690, May 2007. 349.

3. Zahra Batool, Muhammad Usman, Khalid Saleem, M. Abdullah-Al-Wadud Fazal-e-Amin, and Abdulhameed Al-Eliwi. Disease–Disease Association Using Network Modeling: Challenges and Opportunities. Journal of Medical Imaging and Health Informatics, 8(4):627–638, May 2018. 359.

4. Eduardo P. García del Valle, Gerardo Lagunes García Lucía Prieto Santamaría, Massimiliano Zanin, Ernestina Menasalvas Ruiz, and Alejandro Rodríguez-González. Disease networks and their contribution to disease understanding: A review of their evolution, techniques and data sources. Journal of Biomedical Informatics, 94:103206, June 2019. 350.

5. Gerhard H. Scholz and Markolf Hanefeld. Metabolic Vascular Syndrome: New Insights into a Multidimensional Network of Risk Factors and Diseases. Visceral Medicine, 32(5):319–326, 2016. 377.

6. Albert-László Barabási, Natali Gulbahce, and Joseph Loscalzo. Network medicine: a network-based approach to human disease. Nature reviews genetics, 12(1):56–68, 2011.

7. Robert H Eckel, Scott M Grundy, and Paul Z Zimmet. The metabolic syndrome. Lancet, 2005. 376.

8. Ming Lu, Qipeng Zhang, Min Deng, Jing Miao, Yanhong Guo, Wei Gao, and Qinghua Cui. An Analysis of Human MicroRNA and Disease Associations. PLoS ONE, 3(10), October 2008.

9. Silpa Suthram, Joel T. Dudley, Annie P. Chiang, Rong Chen, Trevor J. Hastie, and Atul J. Butte. Network-Based Elucidation of Human Disease Similarities Reveals Common Functional Modules Enriched for Pluripotent Drug Targets. PLoS Computational Biology, 6(2), February 2010.

10. A. Rzhetsky, D. Wajngurt, N. Park, and T. Zheng. Probing genetic overlap among complex human phenotypes. Proceedings of the National Academy of Sciences, 104(28):11694–11699, July 2007. 370.

11. César A. Hidalgo, Nicholas Blumm, Albert-László Barabási, and Nicholas A. Christakis. A Dynamic Network Approach for the Study of Human Phenotypes. PLoS Computational Biology, 5(4):e1000353, April 2009. 371.

12. Yefei Jiang, Shuangge Ma, Ben-Chang Shia, and Tian-Shyug Lee. An Epidemiological Human Disease Network Derived from Disease Co-occurrence in Taiwan. Scientific Reports, 8(91), December 2018. 372.

13. W. Robb MacLellan, Yibin Wang, and Aldons J. Lusis. Systems-based approaches to cardiovascular disease. Nature Reviews Cardiology, 9(3):172–184, March 2012. 397.

14. Gemma M. Kirwan, Diego Diez, Jesper Z. Haeggström, Susumu Goto, and Craig E. Wheelock. Systems Biology Approaches for Investigating the Relationship Between Lipids and Cardiovascular Disease. Current Cardiovascular Risk Reports, 5(1):52–61, February 2011. 396.

15. Anida Sarajlić and Nataša Pržulj. Survey of Network-Based Approaches to Research of Cardiovascular Diseases. BioMed Research International, 2014:1–10, 2014. 398.

16. Babak Ravandi and Arash Ravandi. Network-Based Approach for Modeling and Analyzing Coronary Angiography. 1909.02664 [physics, q-bio, stat], pages 170–181, 2020. 399.

17. Ronald Cornet and Nicolette de Keizer. Forty years of SNOMED: a literature review. BMC Medical Informatics and Decision Making, 8(Suppl 1):S2, October 2008.

18. R. T. Mankowski, S. Yende, and D. C. Angus. Long-term impact of sepsis on cardiovascular health. Intensive Care Medicine, 45(1):78–81, January 2019.

19. Lisa Q. Rong, Antonino Di Franco, and Mario Gaudino. Acute respiratory distress syndrome after cardiac surgery. Journal of Thoracic Disease, 8(10):E1177–E1186, October 2016.

20. R. Scott Stephens, Ashish S. Shah, and Glenn J. R. Whitman. Lung Injury and Acute Respiratory Distress Syndrome After Cardiac Surgery. The Annals of Thoracic Surgery, 95(3):1122–1129, March 2013. Publisher: Elsevier.

21. Jun-Jun Yeh, Cheng-Li Lin, and Chia-Hung Kao. Relationship between pneumonia and cardiovascular diseases: A retrospective cohort study of the general population. European Journal of Internal Medicine, 59:39–45, 2019.

22. Sarmad Said and German T. Hernandez. The link between chronic kidney disease and cardiovascular disease. Journal of Nephropathology, 3(3):99–104, July 2014.

23. Ilan Goldenberg and Arthur J. Moss. Long QT Syndrome. Journal of the American College of Cardiology, 51(24):2291–2300, June 2008.

24. Schwartz Peter J., Crotti Lia, and Insolia Roberto. Long-QT Syndrome. Circulation: Arrhythmia and Electrophysiology, 5(4):868–877, August 2012. Publisher: American Heart Association.

25. El-Sherif Nabil, Chinushi Masaomi, Caref Edward B., and Restivo Mark. Electrophysiological Mechanism of the Characteristic Electrocardiographic Morphology of Torsade de Pointes Tachyarrhythmias in the Long-QT Syndrome. Circulation, 96(12):4392–4399, December 1997. Publisher: American Heart Association.

26. W. Whang, H. M. Julien, L. Higginbotham, A. V. Soto, N. Broodie, J. T. Bigger, H. Garan, M. M. Burg, and K. W. Davidson. Women, but not men, have prolonged QT interval if depressed after an acute coronary syndrome. Europace, 14(2):267–271, February 2012.

27. Ashraf Abugroun, Hussein Daoud, Manar Elhassan, and Dennis Levinson. Coronary Artery Disease Risk Factor Analysis in an Age-Stratified Hospital Population with Systemic Lupus Erythematosus. Journal of the American College of Cardiology, 75(11 Supplement 1):1999, March 2020. Publisher: Journal of the American College of Cardiology Section: Prevention.

28. Ekrem Yasa, Fabrizio Ricci, Martin Magnusson, Richard Sutton, Sabina Gallina, Raffaele De Caterina, Olle Melander, and Artur Fedorowski. Cardiovascular risk after hospitalisation for unexplained syncope and orthostatic hypotension. Heart, 104(6):487–493, March 2018. Publisher: BMJ Publishing Group Ltd and British Cardiovascular Society Section: Cardiac risk factors and prevention.

29. Keld Kjeldsen. Hypokalemia and sudden cardiac death. 15(4):4, 2010.

30. Peter A. McCullough and Norman E. Lepor. The deadly triangle of anemia, renal insufficiency, and cardiovascular disease: implications for prognosis and treatment. Reviews in Cardiovascular Medicine, 6(1):1–10, 2005.

31. Felicita Andreotti, Giulio Coluzzi, Teodosio Pafundi, Teresa Rio, Eliano Pio Navarese, Filippo Crea, Massimo Pistolesi, Attilio Maseri, and Charles H. Hennekens. Anemia contributes to cardiovascular disease through reductions in nitric oxide. Journal of Applied Physiology, 122(2):414–417, February 2017.

32. Fré Bauters, Ernst R. Rietzschel, Katrien B. C. Hertegonne, and Julio A. Chirinos. The Link Between Obstructive Sleep Apnea and Cardiovascular Disease. Current Atherosclerosis Reports, 18(1):1, January 2016.

33. Tuukka Tarvasmäki, Veli-Pekka Harjola, Jukka Tolonen, Krista Siirilä-Waris, Markku S Nieminen, and Johan Lassus. Management of acute heart failure and the effect of systolic blood pressure on the use of intravenous therapies. European Heart Journal. Acute Cardiovascular Care, 2(3):219–225, September 2013.

34. David C Kaelber, Wendy Foster, Jason Gilder, Thomas E Love, and Anil K Jain. Patient characteristics associated with venous thromboembolic events: a cohort study using pooled electronic health record data. Journal of the American Medical Informatics Association, 19(6):965–972, November 2012.

35. Cristian Palmiere and Patrice Mangin. Urea nitrogen, creatinine, and uric acid levels in postmortem serum, vitreous humor, and pericardial fluid. International Journal of Legal Medicine, 129(2):301–305, March 2015.

36. Claire Rimes-Stigare, Bo Ravn, Akil Awad, Klara Torlén, Claes-Roland Martling, Matteo Bottai, Johan Mårtensson, and Max Bell. Creatinine-and Cystatin C-Based Incidence of Chronic Kidney Disease and Acute Kidney Disease in AKI Survivors. Critical Care Research and Practice, 2018:1–8, September 2018.

37. Mathieu Bastian, Sebastien Heymann, and Mathieu Jacomy. Gephi: An Open Source Software for Exploring and Manipulating Networks. In Third International AAAI Conference on Weblogs and Social Media, March 2009.

38. Thomas M. J. Fruchterman and Edward M. Reingold. Graph drawing by force-directed placement. Software: Practice and Experience, 21(11):1129–1164, 1991. _eprint: https://onlinelibrary.wiley.com/doi/pdf/10.1002/spe.4380211102.

39. S. R. Rassekh, M. Lorenzi, L. Lee, S. Devji, M. McBride, and K. Goddard. Reclassification of ICD-9 Codes into Meaningful Categories for Oncology Survivorship Research, 2010.

40. Sameer D. Saini. Cost-effectiveness of Proton Pump Inhibitor Cotherapy in Patients Taking Long-term, Low-Dose Aspirin for Secondary Cardiovascular Prevention. Archives of Internal Medicine, 168(15):1684, August 2008.

41. David A. Peura and C. Mel Wilcox. Aspirin and Proton Pump Inhibitor Combination Therapy for Prevention of Cardiovascular Disease and Barrett’s Esophagus. Postgraduate Medicine, 126(1):87–96, January 2014.

42. George S. Alexopoulos, Jo Anne Sirey, Samprit Banerjee, Dimitris N. Kiosses, Cristina Pollari, Richard S. Novitch, Amanda Artis, and Patrick J. Raue. Two Behavioral Interventions for Patients with Major Depression and Severe COPD. The American Journal of Geriatric Psychiatry, 24(11):964–974, November 2016.

43. Hong Zheng, Peng Zheng, Liangcai Zhao, Jianmin Jia, Shengli Tang, Pengtao Xu, Peng Xie, and Hongchang Gao. Predictive diagnosis of major depression using NMR-based metabolomics and least-squares support vector machine. Clinica Chimica Acta, 464:223–227, January 2017.

