## Supplementary material for "Disease Network Delineates the Disease Progression Profile of Cardiovascular Diseases": Table (1-2), Supplementary Figures (1-6), Supplementary Tables (1-4): Supplementary Materials.docx

Identification of progression rate threshold

To identify the threshold of progression rate, we used a permutation strategy to get progression rate value from randomly selected disease pairs. We sampled 2 diseases without replacement among the diseases with at least 100,000 patients records to build a disease pair. Next, we calculate the PR value of it. Further, we repeated 1,000 times for permutation. The median PR value of these 1,000 values were identified as $PR_{random}$ value of the disease network. Finally, we get $PR_{random} = 2.22$ in general disease network (from age 18 to 90), $PR_{random} = 2.09$ in disease network of young group, $PR_{random} = 2.14$ in disease network of middle-aged group and $PR_{random} = 2.19$ in disease network of the elderly group. Since the diseases we sampled are frequently diagnosed among population, the $PR_{random}$ may be overestimated. Thus, we used $PR = 2.5$ as the threshold of progression rate.
