## Supplementary figures and images for "Disease Network Delineates the Disease Progression Profile of Cardiovascular Diseases"

### Sfig1.pdf

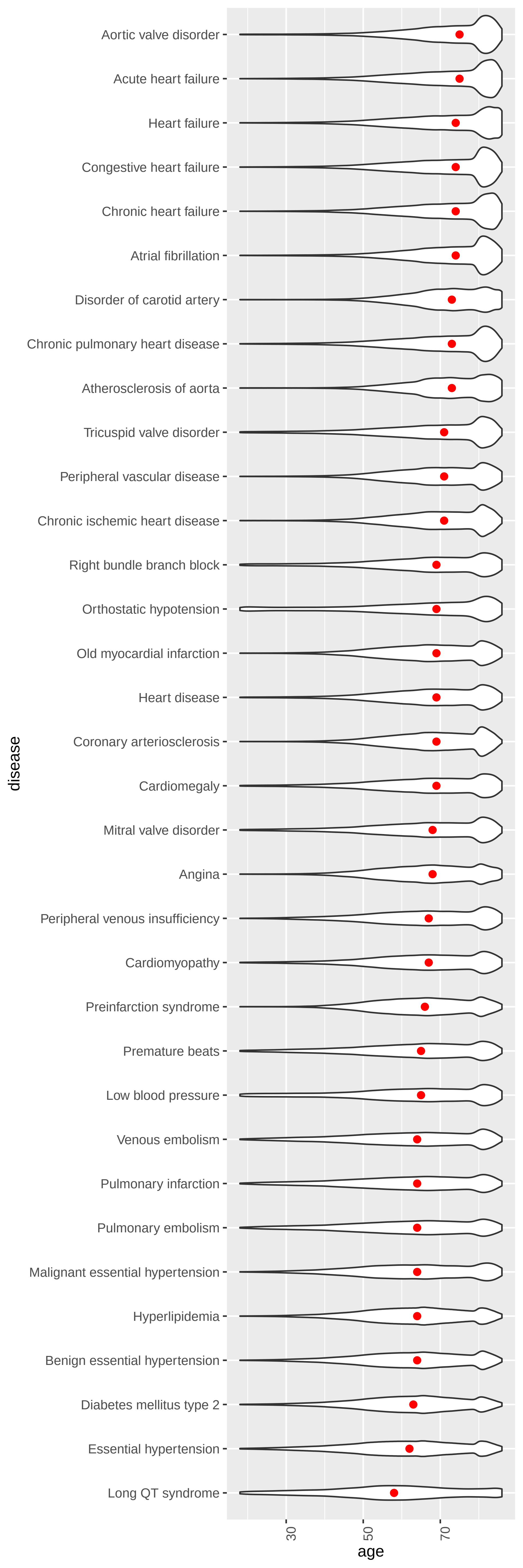

### Sfig2.pdf

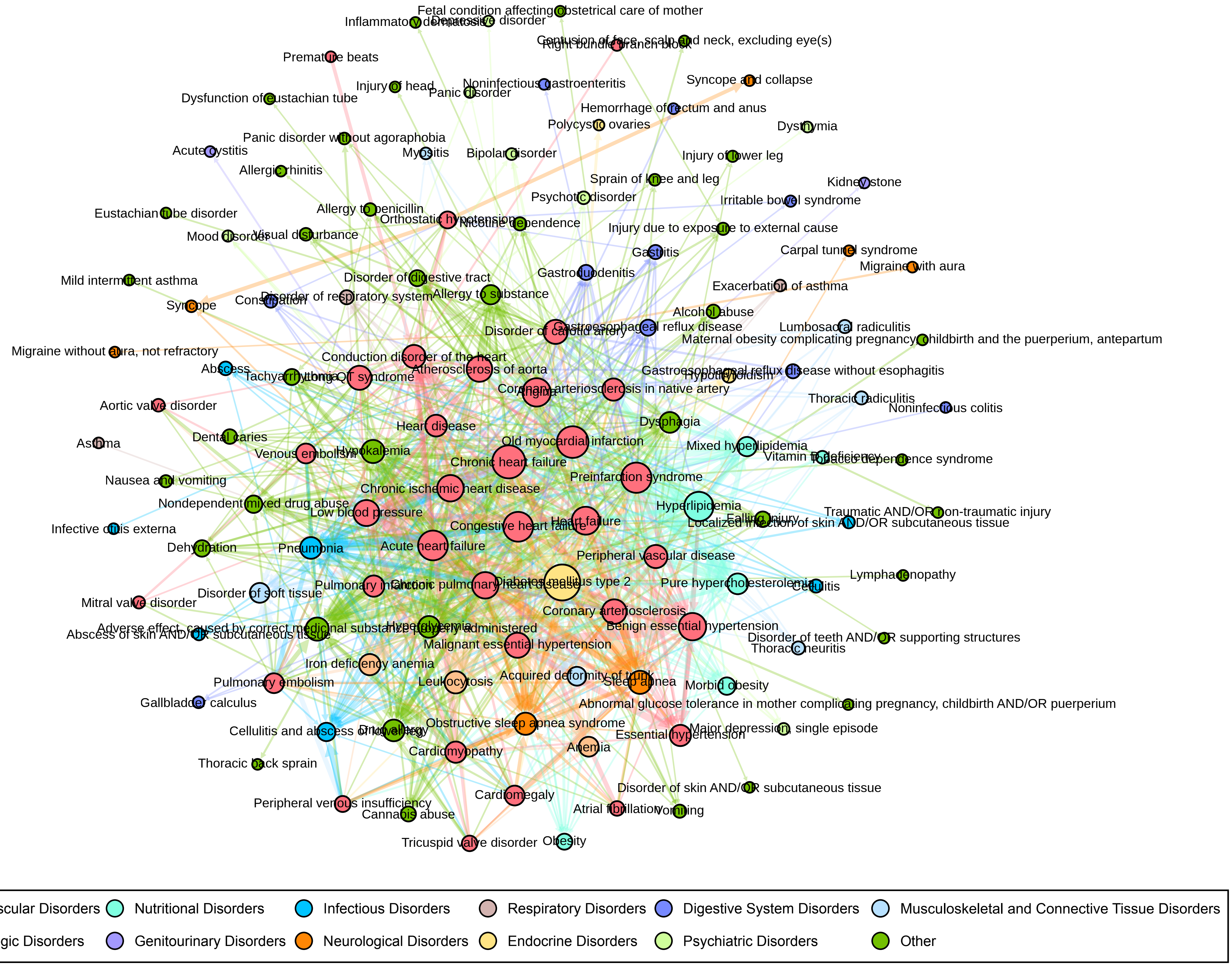

### Sfig3.pdf

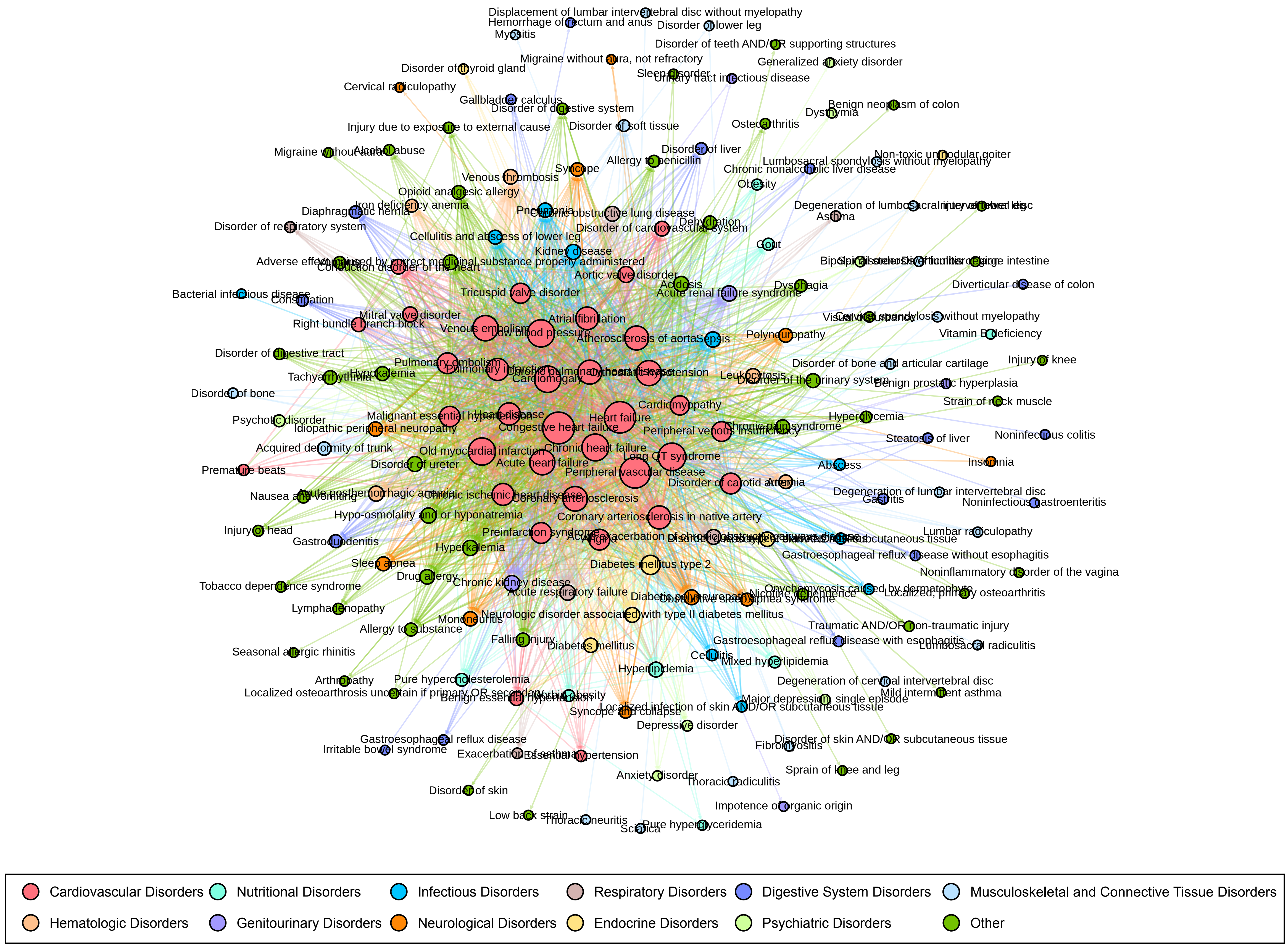

### Sfig4.pdf

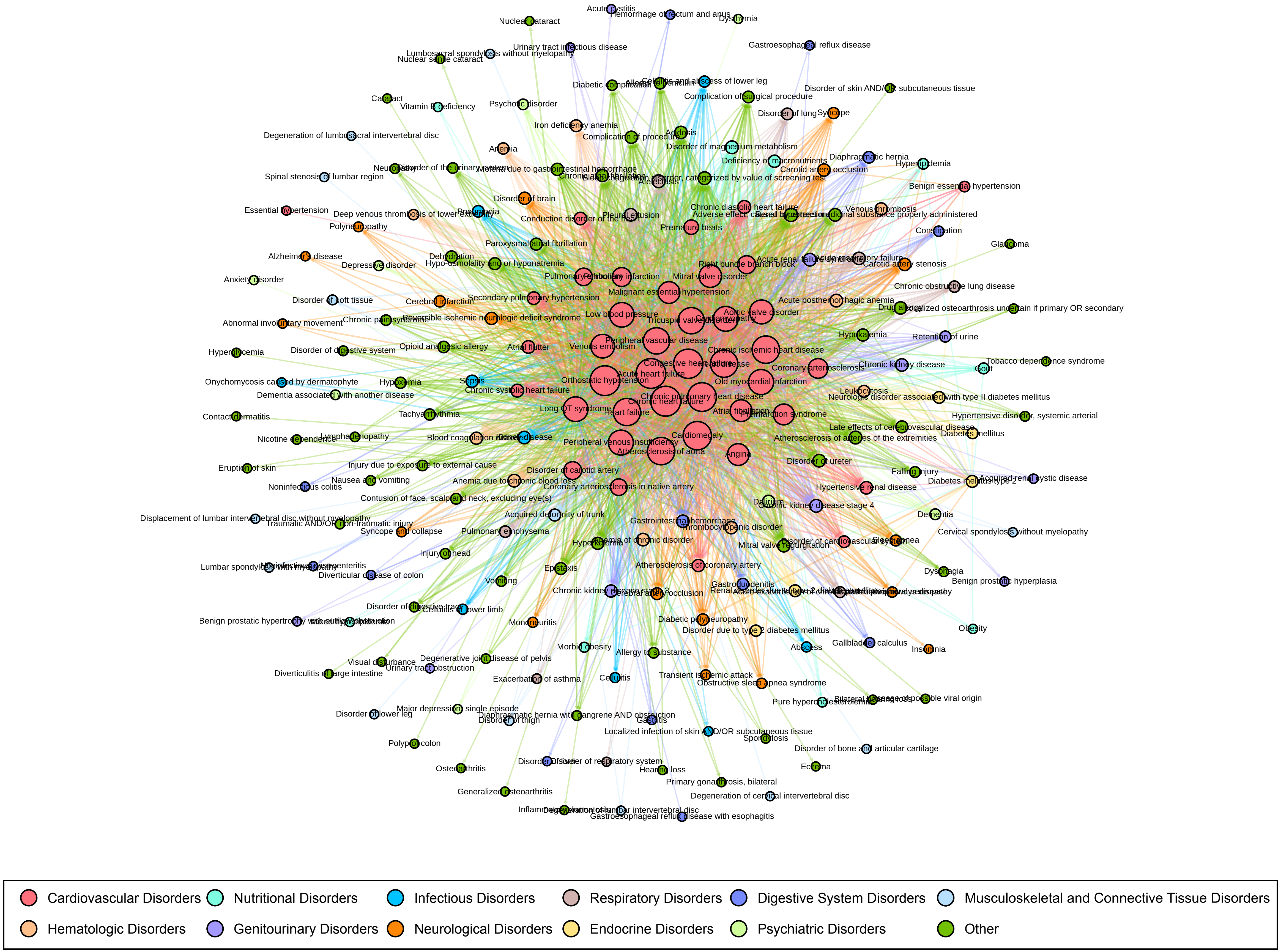

### Sfig5.pdf

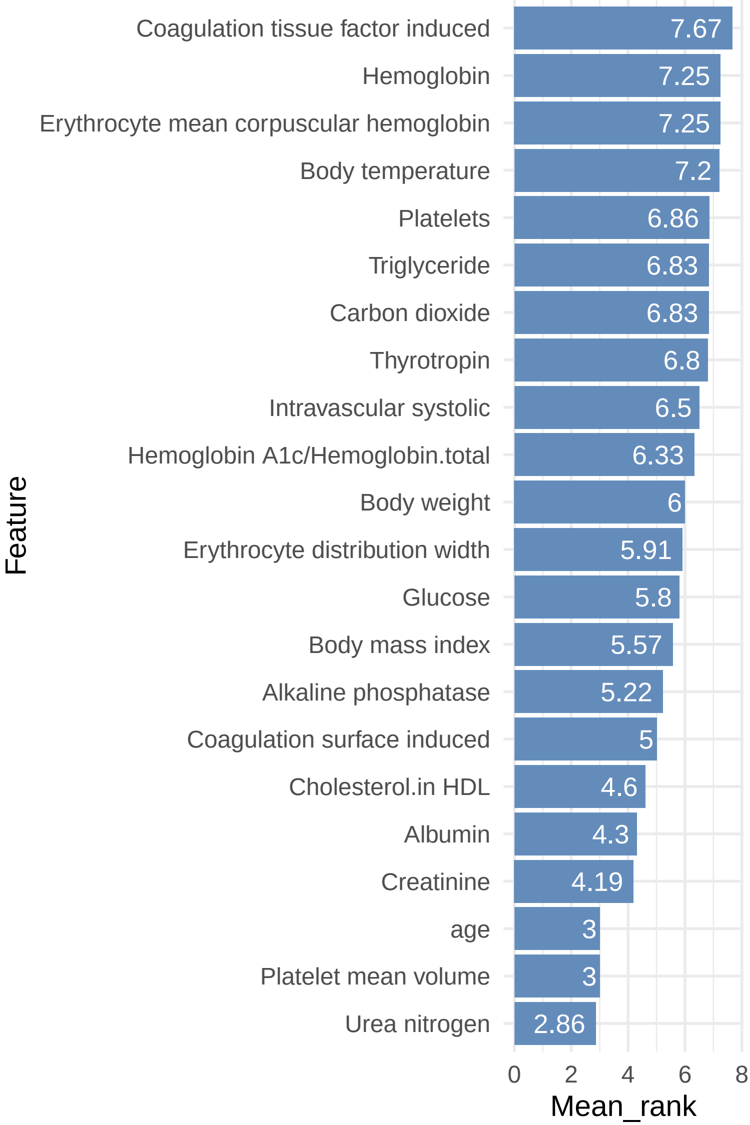

### Sfig6.pdf

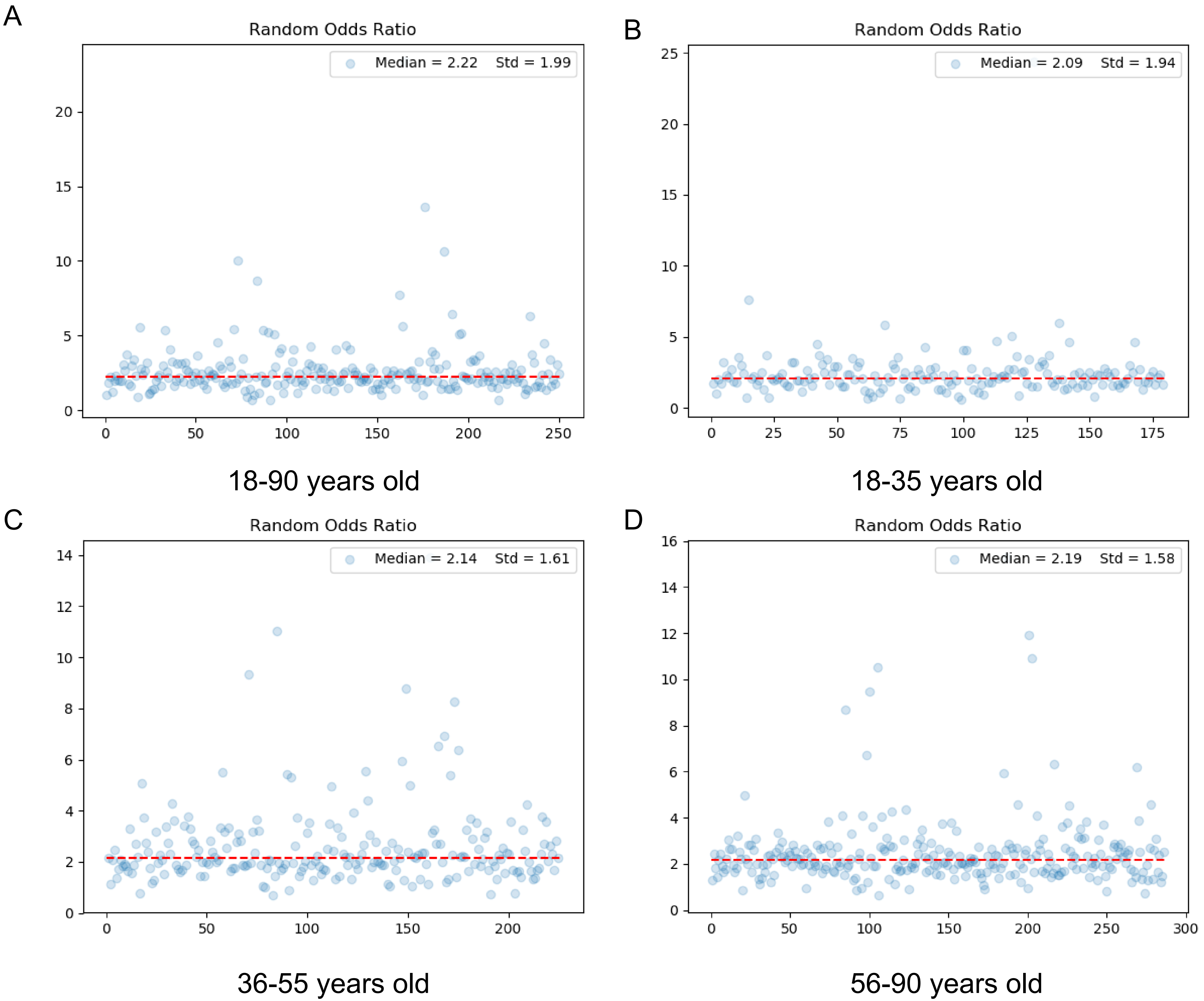
